# Mining of thousands of prokaryotic genomes reveals high abundance of prophage signals

**DOI:** 10.1101/2021.10.20.465230

**Authors:** Gamaliel López-Leal, Laura Carolina Camelo-Valera, Juan Manuel Hurtado-Ramírez, Jérôme Verleyen, Santiago Castillo-Ramírez, Alejandro Reyes-Muñoz

## Abstract

Phages and prophages are one of the principal modulators of microbial populations. However, much of their diversity is still poorly understood. Here, we extracted 33,624 prophages from 13,713 complete prokaryotic genomes in order to explore the prophage diversity and their relationships with their host. Our results reveal that prophages were present in 75% of the genomes studied. In addition, Enterobacterales were significantly enriched in prophages. We also found that pathogens are a significant reservoir of prophages. Finally, we determined that the prophage relatedness and the range of genomic hosts were delimited by the evolutionary relationships of their hosts. On a broader level, we got insights into the prophage population, identified in thousands of publicly available prokaryotic genomes, by comparing the prophage distribution and relatedness between them and their hosts.

## Introduction

Prokaryotic viruses (phages) are considered the most abundant biological entities on the planet [1]. These phages reproduce either through a lytic cycle, like virulent phages, or vertically by integrating into the host genome and taking advantage of its replication cycle as prophages [2]. This integration into the host chromosome can alter the host phenotype by disruption of open reading frames and by altering the expression of flanking genes. Furthermore, this process contributes as a major source of new genes and functions within the bacterial genome [3, 4]. This gene turnover contributes to the fitness and to the appearance of ecologically important bacterial traits such as virulence factors, drug resistance mechanisms [5–7] or phage-derived bacteriocins and tailocins [8, 9]. In this regard, there are some reports that prophages are more frequently present in pathogenic than in non-pathogenic strains [10]. For example, the comparison genomes of laboratory strains from *Escherichia coli* and their pathogenic counterparts revealed that the main differences between these strains were due to the insertion of prophages or other genetic mobile elements [11–13]. Moreover, pathogenicity islands, commonly acquired by horizontal gene transfer are a general mechanism by which many bacteria display a pathogenic phenotype [12]. These observations suggest that the prophages play an important role in the evolution of their hosts [14]. However, these findings come from studies focusing on very particular species.

Until not long ago, the identification of prophage regions had been computationally challenging due to the lack of information about the diversity of phage sequences. However, recently, along with the development of sequencing technologies, the expansion of phage-derived sequence databases has been increasing and, with it, powerful tools for studying phages and prophages [15]. These recent developments have allowed to efficiently identify prophage regions from prokaryotic genomes [16]. Moreover, the advance in sequencing technologies has produced dozens to hundreds of high quality bacterial and archaeal genomes [17, 18], and with this, we have unintentionally sequenced thousands of prophages. Since prophages are part of the host genome, they are *in situ* recovery from their host provides vast information on their diversity and on the relationship with their host. In this sense, few studies have explored the diversity of prophages from genomes and publicly available metagenomic data for elucidating prophage-bacteria relationships [19–21]. However, those studies have analyzed modest numbers of bacterial genomes and have focused mainly on very particular aspects, such as horizontal gene transfer [20], the relationships of the prophage-CRISPR-Cas systems, contribution of host genome size and prophage acquisition and the dominance of commensal lysogens in a particular niche [19]. Nonetheless, no much attention has been paid to other variables such as: i) the phylogenetic relationships of the host and the abundance of the prophages, ii) the presence of prophages in pathogens and non-pathogens and, iii) the host range of prophages and their relatedness in genetic repertoires. Therefore, to analyze these variables and to obtain insights into the knowledge associated with the prophage diversity and their lysogens, we used comparative genomics to characterize the diversity of prophages already identified in over ten thousand prokaryotic genomes.

## Methods

### Prophage prediction

We downloaded all the complete genomic sequences of prokaryotic organisms (13,713 genomes) reported in the RefSeq NCBI database at the end of January 2020. From these, we kept genomes with taxonomic information available at least at the genus level, to avoid potentially misclassification of genomes within the NCBI database [22, 23]. To ensure reliable recovery of prophages, we removed replicons or genomes of less than 10 Kb [15]. Prophage prediction was carried out using VirSorter [15] with default parameters. Prophages from category 1 and category 2 (according to VirSorter) were collected and validated using VIRALVERIFY [24] by removing all prophages predictions that were tagged as plasmids. The resulting prophages were considered as *bona fide* prophages and were kept for downstream analysis. The Wilcoxon test was performed on the distributions of prophages for each taxonomic level (host) in order to evaluate the enrichment of prophages.

### Leave-One-Out approach (LOO)

The Biosample information was obtained from the genomes listed in **Supplementary data 1** using efetch form E-utilities [25]. We collected the metadata of the following sections: general description, isolation source, isolation site, host, environmental medium and sample type. Then, to consider whether a genome corresponds to a pathogenic or non-pathogenic phenotype, we manually checked the information of each section. If they were listed as “pathogen” or “pathogenic” in any of the aforementioned fields, or if the isolation source was from a patient, animal or plant associated with a disease, they were considered pathogens. Otherwise, they were considered as non-pathogenic. Next, in order to determine the contribution of each genus in the whole distribution of the number of prophages for each group (pathogens and non-pathogens), we took out one genus from the at a time from each group of pathogens and non-pathogens, termed the Leave-One-Out (LOO) approach. Statistical test (Wilcoxon test) was carried out with the wilcox.test as implemented in R. We considered an adjusted *P*-value less than or equal to 0.05 as an indicator of statistical significance. The *P*-value correction was performed by the p.adjust function in R using the Bonferroni method.

### Prophage clustering

All the prophages with at least 90% nucleotide similarity over at least 80% of the genome length were clustered using cd-hit [26]. Furthermore, analysis of Average Amino acid Identity (AAI) was carried out based on a pairwise comparison of the dereplicated prophage sequences (clustered prophages) using CompareM (https://github.com/dparks1134/CompareM). We considered relatedness at genus level for those prophages with an AAI value ≥ 80, as previously reported [27, 28]. The prophage relatedness was visualized as an AAI network using Cytoscape v.3.8 [29].

### K-mer usage bias

The k-mer bias measures were obtained for selected bacterial and phages genomes using the k-mer sizes of 1, 2, 3, 4. The calculation of the k-mer usage bias was performed according to the mathematical formula proposed in previous studies [30].

### Tree of life

The Tree of Life (TOL) at order level for Archaea and Bacteria was generated using Lifemap [31]. The TOL was annotated in iTOL [32].

## Results

### Distribution of prophages by host genome size

To explore the presence of prophages in prokaryotic organisms (Bacteria and Archaea), we created a database of 13,713 complete genomes. These genomes had to be assigned to at least genus level in the NCBI (**Supplementary data 1**). We searched for prophage signals using VirSorter [15], resulting in 33,624 prophages. Our results show that lysogens (bacteria with at least one prophage predicted) were more common (75.61%) than non-lysogens (24.38%) (**Supplementary data 2**). Next, we wanted to determine if the presence of prophages was biased to certain bacterial taxa. For this we first ruled out that the number of prophages was not influenced by the host genome size. Some studies have reported that the abundance of prophages positively correlates with the genome size of their host [21]. Here, we found a weak positive correlation (R^2^= 0.32) between (spearman’s *p*-value < 2.2^e-16^) the host genome size and the number of prophages identified (Figure 1). However, we found that genomes with sizes between 4 and 7 Mb were enriched in prophages (Figure 1). These results indicated that the abundance of prophages does not only depend on the host-genome size and could be influenced by other factors.

**Figure 1.**
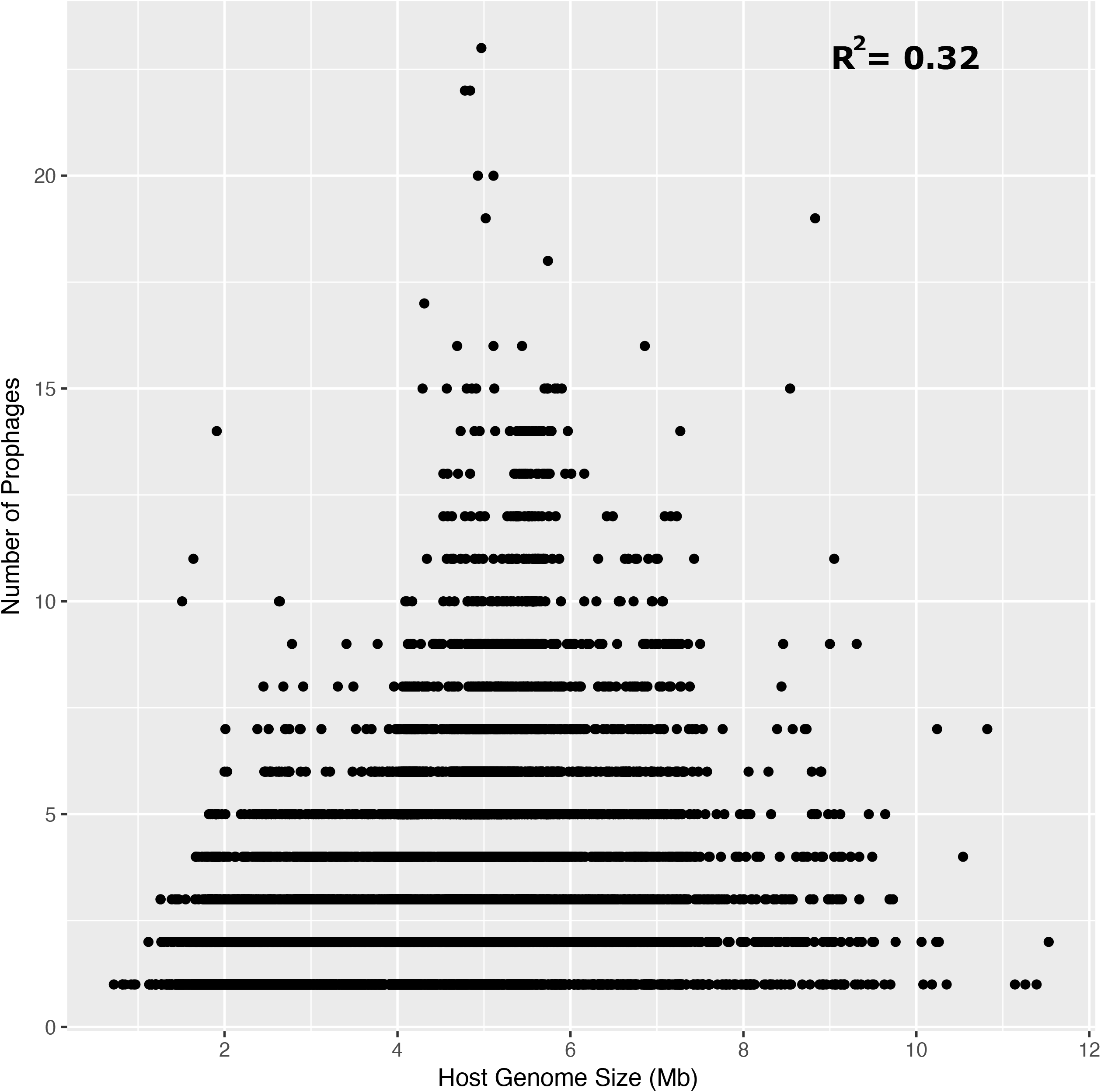
Correlation between the genomes size (Mb) and the number of prophages. Spearman’s rank correlation coefficient is indicated as R value.

### Prophages are enriched in Proteobacteria

Surprisingly, 33,624 prophages were identified from the 10,370 genomes, with a mode and average of 1 and 3.24 prophages, respectively and with a coefficient of variation of 60. We found that the genera *Arsenophonus* and *Plautia*, which belong to the Enterobacterales order, had some of the highest number of prophages, with 23 and 10 detected prophage per genome respectively; however, these genera only had one sequenced representative in our data set. Therefore, to avoid counts of prophages derived from a single representative, we collected all genera that were represented with at least 5 genomes. We found that the members of Proteobacteria phylum showed more prophages (Figure 2). From the lower host taxonomy rank, we found that *Shigella* was the genus with more prophages, with 878 prophages identified in 92 genomes and with a prophage average of 9.54, followed by *Brevibacillus*, *Escherichia* and *Xenorhabdus* with a prophage average of 6.66, 6.43 and 6.14, respectively. Of these, only *Shigella* and *Escherichia* were significantly enriched in prophages (Figure 3A). Moreover, at family level, Morganellaceae, Enterobacteriaceae and Bacillaceae were also significantly abundant in prophages (Figure 3B). Following on this point, only the Enterobacterales order (with a prophage average of 5) was significantly enriched in prophages, and thus, Proteobacteria was the phylum with more abundant prophage signals (Figure 3C, D), despite that Planctomycetales and Entomoplasmatales showed abundant prophages (Figure 2).

**Figure 2.**
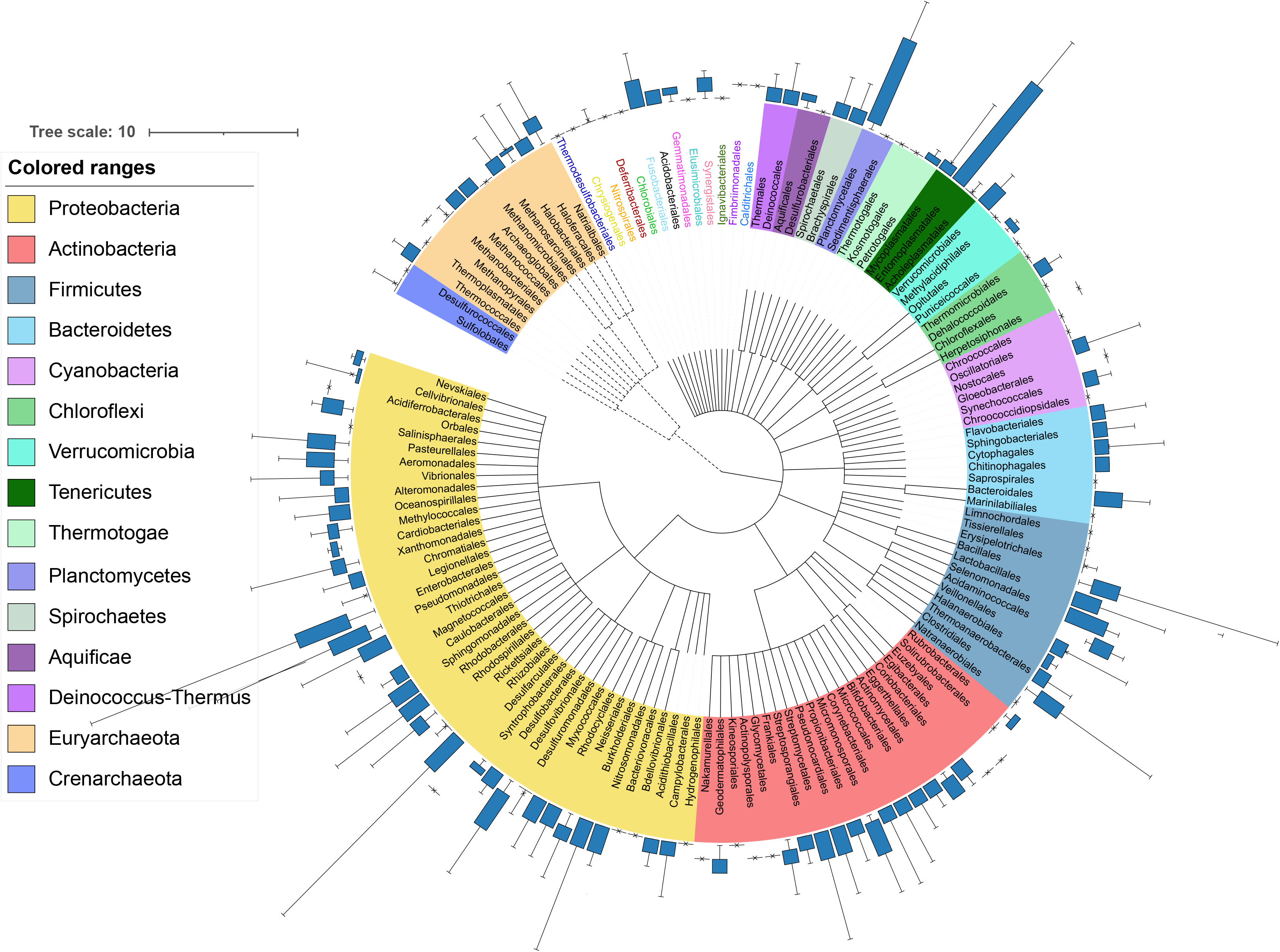
Bacteria and Archaea prophage distribution. The phylogenetic tree was generated using Lifemap at order level. The phylum level is shown with different colors. The Archaea and Bacteria branches are showed in dashed and solid lines, respectively.

**Figure 3.**
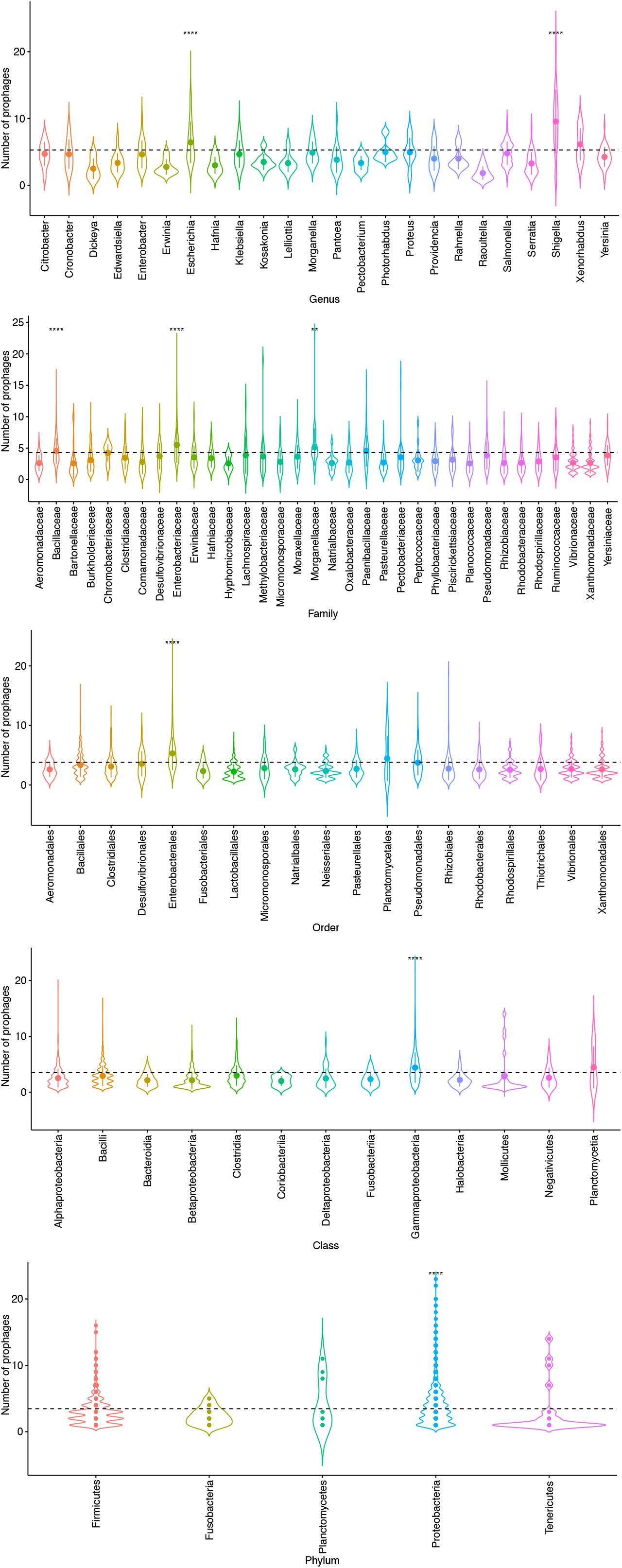
Prophage abundance. Violin plots showing the prophage distributions identified per genome at different taxonomic levels. For all panels, the brackets indicate the significant Wilcoxon tests (p <0.05) of each group (with at least 5 genomes per taxonomic rank) compared against the entire distribution.

It is important to note that the orders with the most genomes were Enterobacterales (2,633), Bacillales (1,419), Lactobacillales (922) and Burkholderiales (916), so to determine if the following orders were enriched in prophages below Enterobacterales, we performed a statistical analysis leaving the Enterobacterales order out. Interestingly, we did not find any significantly enriched order in prophages. These results indicate that the members of the Enterobacterales are significantly enriched in prophage signals (Figure 2). In this regard (without the Enterobacterales group), we found at genus level that the genera *Acinetobacter*, *Bacillus*, *Brevibacillus*, *Lysinibacillus* and *Paenibacillus* were significantly enriched in prophages (**Supplementary data 3**). In summary, these observations suggest that members of Proteobacteria are the only ones enriched in prophages, comparing with other taxonomic groups (Figure 2), suggesting that the abundance of prophages could be associated with their evolutionary history or with the lifestyle of its host. However, some other taxonomic affiliation showed some enrichment of prophages.

### Pathogens display high abundance in prophages

Previous studies have reported that pathogenic strains harbor high abundance of prophages. However, this observation was carried out in few species, such as *Acinetobacter baumannii*, *Escherichia coli* and *Pseudomonas aeruginosa* [11, 12, 33, 34]. Here, we wanted to test if the lifestyle has influence in the accumulation of prophages. First, we randomly collected 100 genomes for those genera that had more than 100 sequenced genomes (to avoid the bias in the number of genomes for each genus, since there are more sequenced genomes for some genera), resulting in 5,728 genomes, and of these, we selected 4,831 genomes which had associated Biosample information. Finally, we created a dataset for pathogenic (n=1,374 genomes with 4,623 prophages) and non-pathogenic (n=3,457 genomes with 9,210 prophages) strains based on their Biosample information (**Supplementary data 4**). We found that the genomes associated with a pathogenic phenotype had a mean and median of 3.36 and 3, respectively, and 2.66 and 2 for non-pathogenic group (**Supplementary data 5**). A first approximation showed a significant abundance in the group of pathogens (*p*-value = 1.99^e-15^). However, in order to determine the contribution of each genus we used a Leave-One-Out (LOO) approach for this data set (see Methods). We found that all genera in the pathogen group were significantly enriched (the 136 genera that make up the pathogen group). On the other hand, 109 genera were shared between the pathogenic and non-pathogenic groups, since most known bacterial pathogens have closely related environmental counterparts [11, 12, 33, 34]. We compared the prophage abundance of the 109 genera between the pathogens and non-pathogens, of these, only *Enterobacter* (*p*-value of 0.008), *Acinetobacter* (*p*-value of 0.038) and *Pseudomonas* (*p*-value of 0.039) were significantly enriched in prophages in the pathogenic isolates compared with their non-pathogenic counterparts (**Supplementary data 6**). Interestingly, these results are in agreement with previous reports that compared few genomes [11, 12, 33, 34].

### Narrow host range delimited by host phylogenetic relationship

One of the main challenges in phage biology is to determine the host range and their genomic diversity [35]. Although there are some reports of phages with a wide host range [36, 37], today there is still a debate about how wide the host range of phages actually is.

Here, to determine the genomic phage-host range, we carried out a clustering over the 33,624 prophages identified, resulting in 22,585 clustered prophages (called Viral Cluster, VC). As we expected, we found that most of the prophages had a narrow genomic host range, with all the members of a given cluster associated to the same host at the genus level (88.4%). However, we found 12 VCs were composed of prophages with different host taxonomic affiliation, of which 6 VCs were identified in Proteobacteria genomes (Table 1). The most discrepant case was VC_94, whose members were found in three *Ralstonia* genomes (*Ralstonia solanacearum*) and once in a *Streptomyces* genome (*Streptomyces spongiicola*). Which is relevant since these species belong to the Proteobacteria and Actinobacteria phyla (Table 1). Interestingly, members from VC_94 showed high identity (>95%) with the previously reported *Ralstonia* RSS-phages types (**Supplementary data 7**) [38–40]. Following this point, we carried out a k-mer bias frequency analysis in order to identify if *Streptomyces spongiicola* is a putative specific host of RSS-phages, since viruses often share higher similarity in k-mer patterns with its host [41, 42]. We found that the k-mer frequencies of dinucleotides and trinucleotides clearly separated the RSS-phages (including the phages from the VC_94) from other *Streptomyces* phages previously reported (**Supplementary data 8**), this suggests that the prophage RSS-type sequence present in *Streptomyces spongiicola* is not associated with an infection process by the RSS-type phage.

**Table 1.** Genomic host range of the viral clusters. The Viral Clusters were assigned by 90% nucleotide similarity and 80% coverage using cd-hit (see Methods). The number of prophage identified in the host genome for specific viral cluster are shown. Level of the genomic host range and the Average Nucleotide Identity (ANI) between host are displayed.

| Viral Cluster (VC) Id | Present in | Host range taxonomic level | ANI between hosts |
| --- | --- | --- | --- |
| vOTU_1233 | 7 *Salmonella* and 2 *Klebsiella* genomes | Genus | 79.49% |
| vOTU_94 | 3 *Ralstonia* and 1 *Streptomyces* genomes | Phylum (Proteobacteria, Actinobacteria) | 71.18% |
| vOTU_1254 | 2 *Odoribacter* and 1 *Parabacteroides* genomes | Family (*Odoribacteraceae*, *Tannerellaceae*) | 73.20% |
| vOTU_1158, vOTU_1237, vOTU_51, vOTU_831 | 1 *Mycobacteroides* and 1 *Mycolicibacterium* genomes | Genus | 83.11% |
| vOTU_1201 | 1 *Mixta* and 1 *Salmonella* genome | Family (*Erwinaceae*, *Enterobacteriaceae*) | 77.11% |
| vOTU_1442 | 1 *Sodalis* and 1 *Citrobacter* genomes | Family (*Pectobacteriaceae*, *Enterobacteriaceae*) | 74.60% |
| vOTU_341, vOTU_781 | 1 *Escherichia* and 1 *Shigella* genome | Genus | 99.95% |

In contrast, for the case of VC_1254 (see Table 1), the prophage identified in *Parabacteroides distasonis* (prophage P1097) showed similar k-mer frequency to the *Odoribacter splanchnicus* species (**Supplementary data 9**).

### Genomic relationships between prophages

We also wanted to determine the relatedness between prophages. For this, we performed an AAI pairwise comparison analysis on the clustered prophages, since AAI was established as a reliable metric to obtain phylogenetic relationship between phages [27, 28]. Here, we retrieved 2,485 prophages that showed >80% of AAI (where phages with >80% of AAI could be associated to the same genus [28]), and most of them were identified within Proteobacteria, Actinobacteria, Bacteroides and Firmicutes; of these, most of the prophages with AAI of >80% were prophages from hosts with the same taxonomic affiliation (Figure 4A). However, Bacteroides and Actinobacteria prophages showed certain degree of relatedness with some Proteobacteria and Firmicutes prophages, suggesting that these prophages have a common genetic pool (Figure 4A).

**Figure 4.**
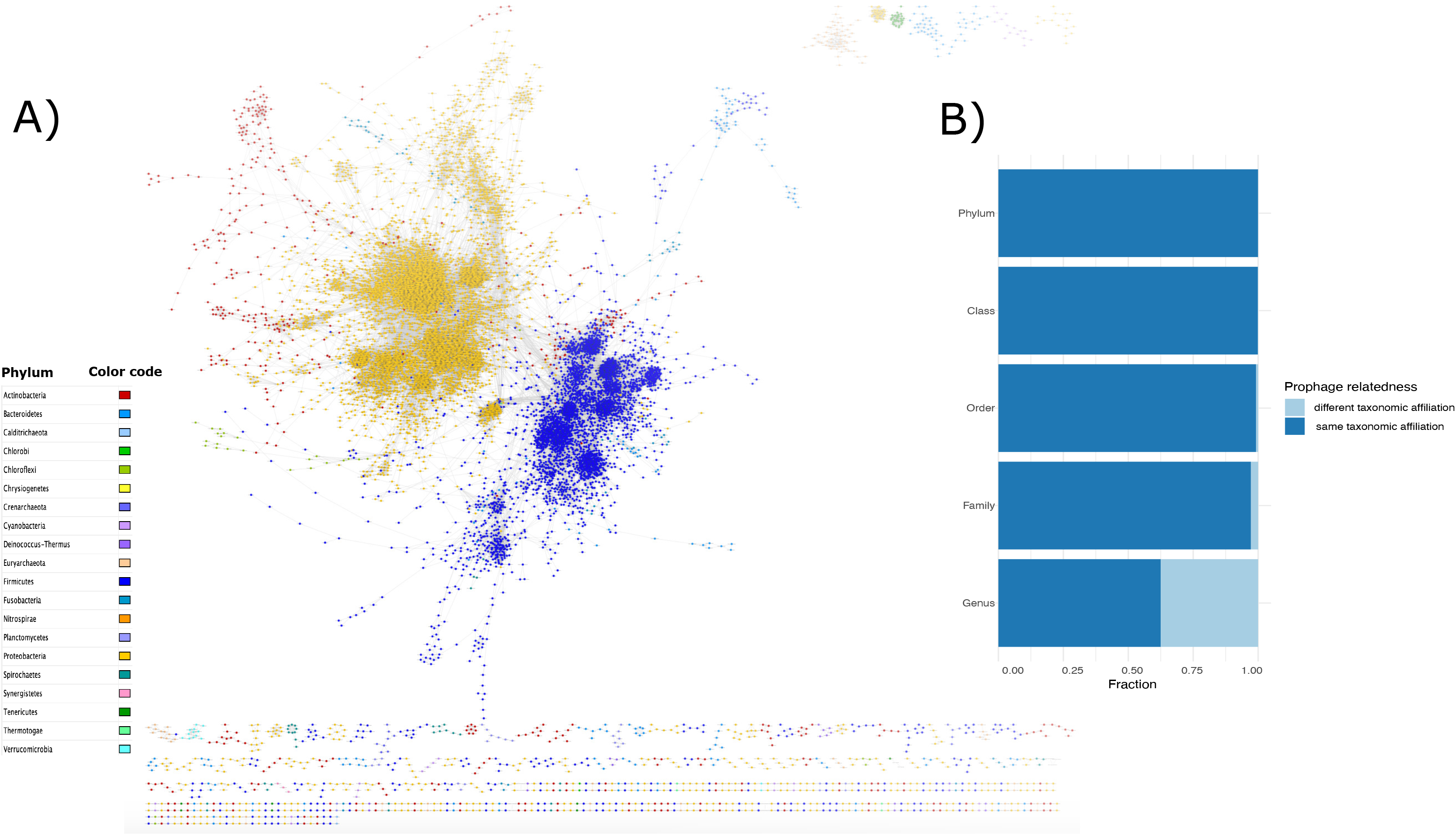
(A) Prophage network generated with prophages with >80 AAI (2,485 prophages), visualization produced with Cytoscape (see methods). Nodes represent prophages and edges represent their weighted pairwise similarities of AAI. Nodes (prophages) are depicted with different colors according to their phylum host. (B) Prophage fraction which shared an AAI value >80 between prophages of different lysogens at all taxonomic levels.

In addition, we determined the prophages relatedness by looking at the fraction of the prophages that had an AAI >80% associated with the same or different host-taxonomic affiliation. We found that at the genus level, the relatedness between prophages from hosts with the same taxonomic affiliation was more common (Figure 4B). However, the relationship between prophages of different lysogens (phylogenetically distant) at the genus level was higher (0.37) and decreased as the phylogenetic relationships of the lysogens were more distant (phylum 0.0002). These results suggest that prophages could preferentially change at genus level or come from a common gene pool, which would cause recombination events between prophages and phages (or other prophages) [43].

## Discussion

To the best of our knowledge, this study comprises the most extensive analysis of prophages and describes the diversity of 33,624 prophages found in 13,713 prokaryotic complete genomes. In contrast to previous studies [21], our results indicate that there is a week correlation between the host genome size and prophage abundance. In this regard, Touchon and colleagues could determine strong positive correlation between the prophage abundance and the host-genome size using 2,110 genomes [21]. This correlation was observed only for genomes with a size up to 6 Mb. Moreover, they selected as *bona fide* those phages with a size above 30Kb. Here, we found that the prophages were enriched in genomes between 4 and 7 Mb from a data set of 14,055 genomes, and the prophages abundance was in accordance with the taxonomy of the host rather than the genome size. This result is consistent with a previous study [19], indicating that temperate phage features have developed over a long phylogenetic timescale. In addition, the prevalence of the lysogens identified concurs to the prevalence of lysogens analyzed in gut metagenomics samples [19], indicating that our observations can also be obtained using different data sets.

We found that the Proteobacteria phylum, as well as the order of the Enterobacterales and most of its affiliated genera, were enriched in prophages, suggesting that the abundance of prophage signals could be associated with the taxonomy of the host [19]. Interestingly, most of the genera enriched in prophages have been placed at the top of the 2017 World Health Organization Priority List for Research and Discovery of New Antibiotics [44], such as: *Shigella*, *Escherichia*, *Acinetobacter*, *Pseudomonas*, etc. (Figure 3 and **Supplementary data 3**), due to their relevance to public health. Therefore, we consider that our results provide evidence to put more effort into studying the phage-bacteria relationships in these genera. Some reports have determined that prophages play a major role in the pathogenicity of their host by the acquisition of ARG and toxins [6, 11–13]. However, this observation has been reported in Bacteria rather than Archaea. Although Archaea prophages have also been studied, Archaea pathogens have not yet been reported. [45].

In this regard, we found that prophages are more abundant in bacteria associated with pathogenic phenotypes. Interestingly, these results agree with previous reports, where clinical isolates of *Acinetobacter baumannii* and *Pseudomonas aeruginosa* [33, 34] showed higher abundance of prophages than environmental isolates. Our results indicated that the *Acinetobacter* and *Pseudomonas* genera were enriched in prophages when these species displayed pathogenic phenotypes (Supplementary data 5). Therefore, this suggests that prophages could play an important role in the appearance of pathogenic phenotypes for some species, since in recent years it has been reported that prophages in *Acinetobacter*, *Pseudomonas*, *Escherichia* and *Shigella* are involved in horizontal transfer of toxin and antibiotic resistance genes [6, 46, 47].

One characteristic of viruses is that they are highly diverse and have multiple phylogenetic origins [48]. Recently, new tools have been developed to determine a good classification of viruses and to infer properly their phylogenetic relationships [28]. In this study, we used two approaches (nucleotide and amino acid approaches) to determine the prophages-relationship, as previously was reported [28]. As expected, most of the prophages were singletons (genome specific); however, we found some VC across different taxonomic groups (Table 1).

Although there are very few studies of lytic phages capable of infecting bacteria of different phyla (because lytic phages usually have a broader host range) [36, 37], there are no reports of prophages that have this characteristic. The case of the RSS-phages that we found in some genomes of *Ralstonia solanacearum* and *Streptomyces spongiicola* HNM0071 is undoubtedly a fascinating case. Nonetheless, the k-mer bias analysis revealed that the RSS-phage found in *Streptomyces spongiicola* HNM0071 could not be associated with an infection process, so we cannot rule out the possibility of contamination until this observation is experimentally proven, although the *Streptomyces spongiicola* HNM0071 genome is of high-quality.

Following this point, we wanted to determine whether temperate phages are closely related to each other (using AAI) and if highly related phages had the possibility of jumping to new hosts (infected by related phages), due to sharing a common gene pool. For this, previous studies have reported AAI values are a good metric to stablish phylogenetic relationships between phages [28]. Our results showed that most of the prophages with high similarity are limited to the evolutionary (taxonomic) scale of their hosts (Figure 4B). Moreover, our results are consistent with other reports where they determined that the gene turnover of phage-host and host-phage relationships is generated principally at the genus and species level and delimited by the evolutionary relationships of their hosts [20, 37]. Interestingly, these studies reported that the gene turnover occurs with high frequency among the Enterobacterales prophages [20]. These observations, together with our results (where Enterobacterales are abundant in prophages), suggest that Enterobacterales could be a hot spot where prophages are the main mechanism that improves the plasticity of the genome of these species.

In addition, it is known that Proteobacteria are the most diverse phylum [49], and in the last decades, a rapid diversification of toxin genes and antibiotic resistance has been observed within the Enterobacteriaceae members [50]. Therefore, future analyzes are needed to test whether phages are largely responsible for such diversification.

Finally, our findings expanded the knowledge of lysogenic interactions between prokaryotes and their prophages. We consider that these associations help to explain the relationships of prophages and their hosts in certain clades; however, more studies are needed to address the degradation processes [43] of prophages and their contribution in the bacterial fitness and the gene turnover.

## Acknowledgements

GLL received a postdoctoral fellowship (2019-000012-01EXTV-00488) from CONACyT. We are thankful to Alfredo Hernández-Alvarez and Víctor Del Moral-Chávez for technical support. GLL thanks to Juan Sebastian Andrade Martinez, Luis Alberto Chica Cárdenas, Laura Milena Forero Junco, Ruth Hernandez Reyes, Leonardo Moreno Gallego, Guillermo Rangel and Laura Avellaneda Franco, members of the Viromics group within the Max Planck Tandem Group in Computational Biology, for the general discussion of the results presented in this manuscript.

## Supplementary figures legends

**Supplementary data 1.** List of genomes used.

**Supplementary data 2.** Prevalence of lysogens and no lysogens per genomes completeness (n= 13,717).

**Supplementary data 3.** Prophage abundance. Violin plots showing the prophage distributions identified per genome at genus level. For all panels, brackets indicate Wilcoxon tests significant (p < 0.05) of each genus (without the Enterobacterales and their affiliated taxonomy ranks) comparisons against the rest.

**Supplementary data 4.** Genomes list collected based on their Biosample information.

**Supplementary data 5.** Boxplots showing the prophage abundance distribution per genome in pathogens (n=1,375 genomes) and non-pathogens (n=3,457).

**Supplementary data 6.** Number of prophages identified in the retrieved genera for the pathogen and non-pathogen group. Standard deviation is shown in the SD column. Statistical test (Wilcoxon test) *p*-values and the *p*-value correction (Bonferroni method) are shown.

**Supplementary data 7.** Genomic synteny comparisons with Easyfig and the BLASTn algorithm. Prophages identified in this study are shown in bold. Arrows represent the locations of coding sequences and shaded lines reflect the degree of homology between pairs of phages. Protein’s functions are displayed in the correspond colors.

**Supplementary data 8.** Heatmap of the K-mer bias frequency (log values) of di and tri-nucleotides are shown. High and low K-mer frequency are displayed in red and blue, respectively. Phages and prophages genomes are shown in red. Bacterial genomes are shown in black.

**Supplementary data 9.** Heatmap of the K-mer bias frequency (log values) of di and tri-nucleotides are shown. High and low K-mer frequency are displayed in red and blue, respectively. Prophages genomes from VC_1254 are shown in red. Bacterial host genomes are shown in black.

